# Projected changes of alpine grassland carbon dynamics in response to climate change and elevated CO_2_ concentrations under Representative Concentration Pathways (RCP) scenarios

**DOI:** 10.1101/595926

**Authors:** Pengfei Han, Xiaohui Lin, Wen Zhang, Guocheng Wang

## Abstract

The Tibetan Plateau is an important component of the global carbon cycle due to the large permafrost carbon pool and its vulnerability to climate warming. The Tibetan Plateau has experienced a noticeable warming over the past few decades and is projected to continue warming in the future. However, the direction and magnitude of carbon fluxes responses to climate change and elevated CO_2_ concentration under Representative Concentration Pathways (RCP) scenarios in the Tibetan Plateau grassland are poorly known. Here, we used a calibrated and validated biogeochemistry model, CENTURY, to quantify the contributions of climate change and elevated CO_2_ on the future carbon budget in the alpine grassland under three RCP scenarios. Though the Tibetan Plateau grassland was projected a net carbon sink of 16 ~ 25 Tg C yr^-1^ in the 21st century, the capacity of carbon sequestration was predicted to decrease gradually because climate-driven increases in heterotrophic respiration (Rh) (with linear slopes 0.49 ~ 1.62 g C m^-2^ yr^-1^) was greater than the net primary production (NPP) (0.35 ~ 1.52 g C m^-2^ yr^-1^). However, the elevated CO_2_ contributed more to plant growth (1.9% ~ 7.3%) than decomposition (1.7% ~ 6.1%), which could offset the warming-induced carbon loss. The interannual and decadal-scale dynamics of the carbon fluxes in the alpine grassland were primarily controlled by temperature, while the role of precipitation became increasingly important in modulating carbon cycle. The strengthened correlation between precipitation and carbon budget suggested that further research should consider the performance of precipitation in evaluating carbon dynamics in a warmer climate scenario.

## 1. Introduction

The distinctive geographic environment of the world highest plateau—Tibetan Plateau (with an average elevation of over 4000 m above sea level) makes its carbon cycle be strongly sensitive to climate variation and environmental change [1-3]. The Plateau covers a vast area of alpine vegetation and has a large reservoir of organic carbon in the permafrost soil, and it has been reported as a significant terrestrial carbon sink for recent decades due largely to warming and CO_2_ fertilization [3-5]. The Plateau has undergone significant warming during the last decades. Increasing temperatures have been reported to prolong growing season lengths [6], metabolic rates, and in the productivity and distribution [7] of vegetation at the Plateau and high latitude areas [8-10]. Furthermore, the fate of soil organic carbon (SOC) is controlled by the complex processes involving accumulation of carbon input by plant production and loss through microbial decomposition. Warming may also suppress plant growth by increasing evapotranspiration [11] and induce the soil carbon loss through enhanced Rh [12]. Thus the carbon fluxes can be amplified by increased carbon emissions from thawing permafrost [13,14]. Schaphoff et al. [15] revealed that warming contributed to the net uptake of carbon in the permafrost zone in previous decades because carbon uptake by vegetation increased at a faster rate than that released from soil. Furthermore, increasing the atmospheric CO_2_ concentrations tends to stimulate photosynthesis and reduce water loss [16,17], which is likely to offset the adverse effects of climate change.

The temperature on the Tibetan Plateau increased with a linear trend of 0.2 °C/decade during the past five decades [18,19] and is projected to continue warming in the future under different representative concentration pathways (RCPs) [20,21]. Although there has been several studies on the carbon cycle and climate change associated with the elevated CO_2_ on the Tibetan Plateau for historical research and sensitivity analysis [2,3,20,22,23], there is still a lack of studies on the carbon dynamic projections under different RCPs on the Plateau. Temperature has been reported as the critical determinant of carbon exchange in alpine ecosystems [24,25]. The warming had prolonged the alpine plant phenology [26-28] and promoted the vegetation productivity [3,29], and consequently enhanced carbon inputs into the soil, offseting the carbon release from thawing permafrost in the Tibetan Plateau [4]. The vegetation production on Tibetan Plateau was projected to increase under future climate scenarios [7,30]. The results of field experiments, however, have shown some opposite responses of vegetation productivity to warming, which varied by regions [31-33]. Ganjurjav et al. [34] found that the aboveground net primary production (NPP) was significantly stimulated in an alpine meadow under experimental warming, whereas it decreased in an alpine steppe due to warming-induced drought. Furthermore, there are situations that the climatic warming stimulated the ecosystem respiration more than NPP, which potentially led to loss of soil carbon in the Tibetan Plateau [18,22,30]. This discrepant responses of NPP and heterotrophic respiration (Rh) to the climatic warming in the Tibetan Plateau might be quite different from that in the northern high-latitude permafrost zones [15], where warming increased NPP more than Rh. In previous historical period modeling studies [3,5,35], a net carbon sink has been estimated for the Tibetan Plateau grassland for the 20th century, albeit with variability in absolute value of 11.8 ~ 29.8 Tg C yr^-1^. Based on a field experiment in the Tibetan Plateau alpine meadow, Zhu et al. [36] found that warming caused a seasonal shift in ecosystem carbon exchange but had little impact on net carbon uptake. However, the direction and magnitude of carbon fluxes responses to climate change and elevated CO_2_ concentration under future climate scenarios in the Tibetan Plateau grassland remains uncertain. Thus, quantifying the influence of the future climatic change on the carbon exchange in alpine ecosystems in the Tibetan Plateau will promote our understanding of the mechanisms in the climate-carbon cycle feedbacks.

In this study, we used an onsite calibrated and validated process-based model, CENTURY, to explore the responses of three important carbon cycle indicators, namely NPP, Rh and net ecosystem production (NEP), to future climate change projections on alpine grasslands of the Tibetan Plateau. The representative concentration pathways (RCP) scenarios of the climate change projections under the framework of the Coupled Model Intercomparison Project Phase 5 (CMIP5) vary in their total radiative forcing increase by 2100 [37,38]. Three different RCP scenarios (i.e., RCP2.6, RCP4.5, and RCP8.5) were applied in this study. The objectives of this paper are: (1) to predict the trends and differences in the ecosystem carbon cycle components in alpine grasslands under RCP scenarios during the 21st century; and (2) to differentiate the contributions of the changing climate variables (temperature and precipitation) and elevated CO_2_ on the future carbon budget on the Tibetan Plateau grasslands.

## 2. Materials and methods

The CENTURY model was originally developed for the U.S. Great Plains grassland ecosystem [39] and has been successfully adapted to simulate carbon fluxes under climate change across a wide latitudinal gradient, from tropical to temperate to high-latitude ecosystems [40-44]. A detailed description of the CENTURY model has been presented by [39,41]. The observed data for parameterization of the CENTURY model included the nutrient contents in vegetation and soil that were derived from the Haibei research station of the Chinese Ecosystem Research Network (CERN), Chinese Academy of Sciencies. The performance of the CENTURY model on the Tibetan Plateau grassland has been evaluated by the observed net ecosystem exchange (NEE), ecosystem respiration (Re), gross primary production (GPP) (2003 ~ 2005) obtained from the Haibei flux tower observations and aboveground biomass (AGC) (2000 ~ 2012) obtained from the field sampling at Haibei research station [23]. In addition, the Re was further separated into autotrophic respiration (Ra) and heterotrophic respiration (Rh) according to the study of [45], and the net primary production (NPP) was then obtained by subtracting Ra from GPP. The site-level verification from our previous study showed that the parameterized CENTURY model was able to capture the carbon fluxes of alpine grassland (R^2^ > 0.80, Fig 1). The simulated monthly AGC exhibited a good agreement with the observed data with the slope close to 1, and the intercept and the root mean square error (RMSE) were less than 25 g C m^-2^. Compared to the eddy-covariance flux data, the simulated monthly NPP and Rh were comparable to the observation with slopes equal to 0.77 and 1.10, and the estimated errors of RMSE were 29.06 g C m^-2^ and 13.24 g C m^-2^, respectively. The simulated NEE was also agreed well with the field observation (R^2^ = 0.88, RMSE = 17.94), while the seasonal amplitude of the simulated NEE was slightly higher than that from the flux tower data with a slope of 0.64) [23]. We acknowledged that the model parameterization was dependent on data from Haibei station, which could cause a certain degree of uncertainty. This was due largely to the fact that most of the existed observations on the Plateau were not long-term experiments and lacked necessary information for model calibration. At the regional scale, our previous historical simulated results were comparable to other studies, with a mean NPP and NEP of 259 g C m^-2^ yr^-1^ and 10 g C m^-2^ yr^-1^ from 1981 to 2010, respectively [23], which was within the range of 120 ~ 340 g C m^-2^ yr^-1^ and 8 ~ 24 g C m^-2^ yr^-1^ reported by [3,5,46]. The spin up procedure for the CENTURY model simulation followed the published literature [47,48].

**Fig 1.**
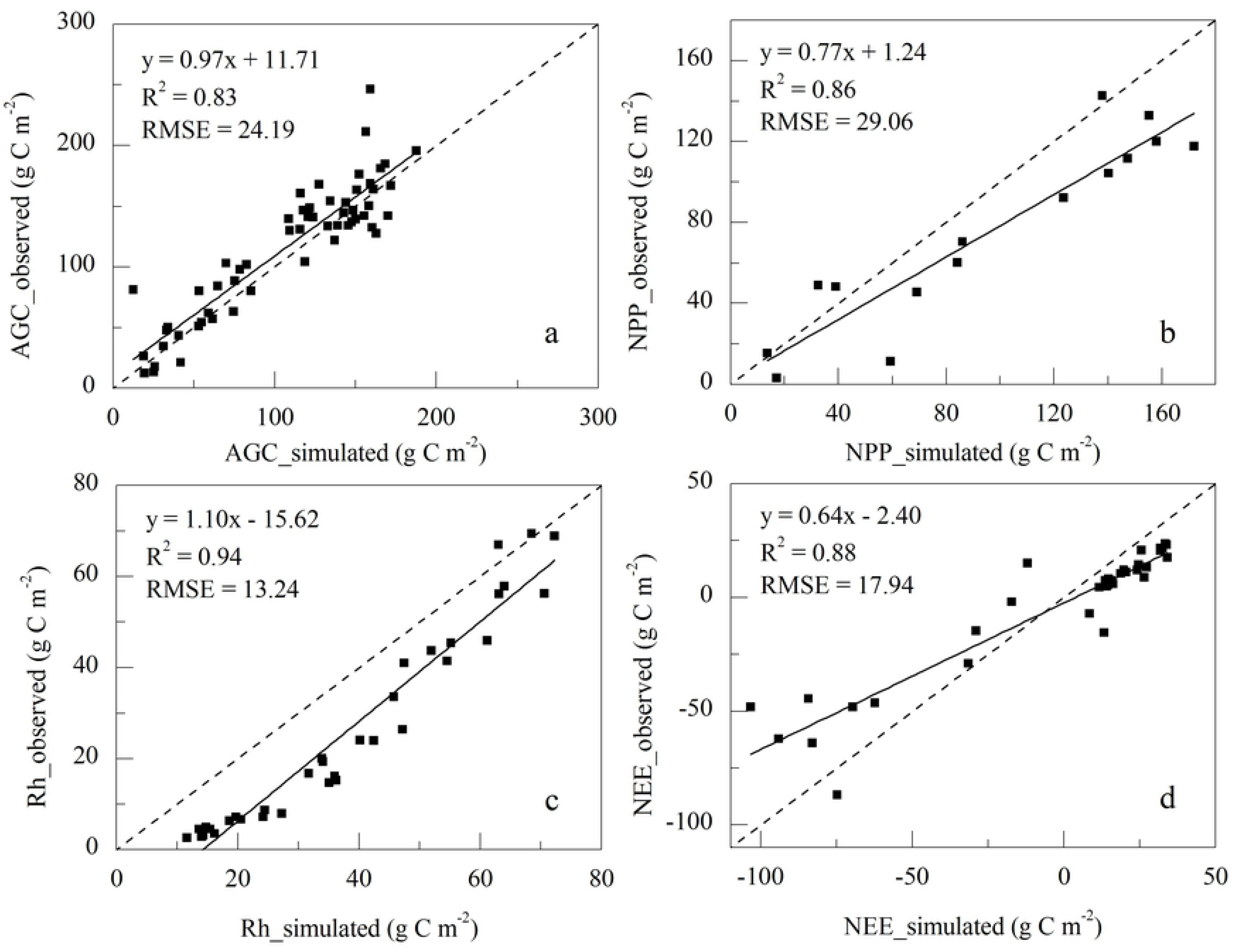
Comparisons of observed and modeled aboveground biomass (AGC, a), and net primary production (NPP, b), heterotrophic respiration (Rh, c), and net ecosystem production (NEE, d) at the Haibei research station of the Tibetan Plateau.

The simulation began with an equilibrium run to generate an initial condition using long-term average climate data for the period from 1901~1930, which was then followed by a spin up (1980 ~ 2005) to eliminate system fluctuations and to steady the transient simulation. The climate input data for the history period of 1901 ~ 2005 was obtained from the Climatic Research Unit (CRU) Time-Series (TS) version 3.23 (TS 3.23), School of Environmental Sciences, University of East Anglia, United Kingdom [49,50]. The CENTURY model was then forced with outputs from the Coordinated Regional Downscaling Experiment (CORDEX), including monthly maximum and minimum temperature and precipitation under three RCPs scenarios (RCP2.6, RCP4.5 and RCP8.5). The CORDEX data used in this study were dynamical downscaled at a high spatial resolution of 0.5° with two regional climate models (RCMs) of RCA4 and REMO2009 driven by ICHEC-EC-EARTH and MPI-M-MPI-ESM-LR global models based on the Coupled Model Intercomparison Project Phase 5 (CMIP5) simulations [51]. The calibrated parameters for the regional simulation were obtained from our previous studies on the Tibetan Plateau grasslands [23].

## 3. Results

### 3.1 Temporal dynamics of the carbon budget on the Tibetan Plateau grasslands

Interannual variations in the grassland carbon budget on the Tibetan Plateau in response to changes in climate and atmospheric CO_2_ over the 21st century were simulated. The CENTURY model predicted that the grass NPP would range from 308 ± 13 g C m^-2^ yr^-1^ to 495 ± 21 g C m^-2^ yr^-1^, with a multiyear mean NPP of 357 ± 18 g C m^-2^ yr^-1^ for RCP2.6, 375 ± 20 g C m^-2^ yr^-1^ for RCP4.5, and 408 ± 26 g C m^-2^ yr^-1^ for RCP8.5 on the Tibetan Plateau, respectively (Fig 2a). The interannual variation in the grass NPP was projected to increase to different degrees under the three RCP scenarios over the period 2006 ~ 2100, with linear slopes of 0.35 g C m^-2^ yr^-1^ (*P*<0.01), 0.85 g C m^-2^ yr^-1^ (*P*<0.01), and 1.52 g C m^-2^ yr^-1^ (*P*<0.01) for RCP2.6, RCP4.5, and RCP8.5, respectively. The temporal dynamics of Rh were consistent with the NPP trends in all scenarios, with mean absolute values of 346 ± 9 g C m^-2^ yr^-1^, 360 ± 12 g C m^-2^ yr^-1^, and 390 ± 19 g C m^-2^ yr^-1^, whereas the magnitude of the variation rates of Rh were all greater than that of NPP, with slopes of 0.49 g C m^-2^ yr^-1^ (*P*<0.01), 0.91 g C m^-2^ yr^-1^ (*P*<0.01), and 1.62 g C m^-2^ yr^-1^ (*P*<0.01) for RCP2.6, RCP4.5, and RCP8.5, respectively (Fig 2b). These higher rates of Rh increase in all the three scenarios may suggest that Rh was more sensitive to climate change than NPP. On average, the alpine grasslands of the Tibetan Plateau behaved as a carbon sink, with a simulated NEP on a range of 11 ± 16 g C m^-2^ yr^-1^, 15 ± 16 g C m^-2^ yr^-1^, and 18 ± 16 g C m^-2^ yr^-1^ for RCP2.6, RCP4.5, and RCP8.5, respectively, for the period from 2006~2100 (Fig 2c). However, the temporal dynamics of the NEP indicated relatively less variability than NPP and Rh under the three RCP scenarios, and decreased continually with the slopes of −0.14 g C m^-2^ yr^-1^ (*P*<0.01), −0.07 g C m^-2^ yr^-1^ (*P*=0.13), and −0.10 g C m^-2^ yr^-1^ (*P*<0.05) for RCP2.6, RCP4.5, and RCP8.5, respectively. This indicated that the potential capacity for carbon sequestration on the Tibetan Plateau grasslands would gradually decrease under future climate change, because the magnitude of the variation rate of Rh was greater than that of NPP.

**Fig 2.**
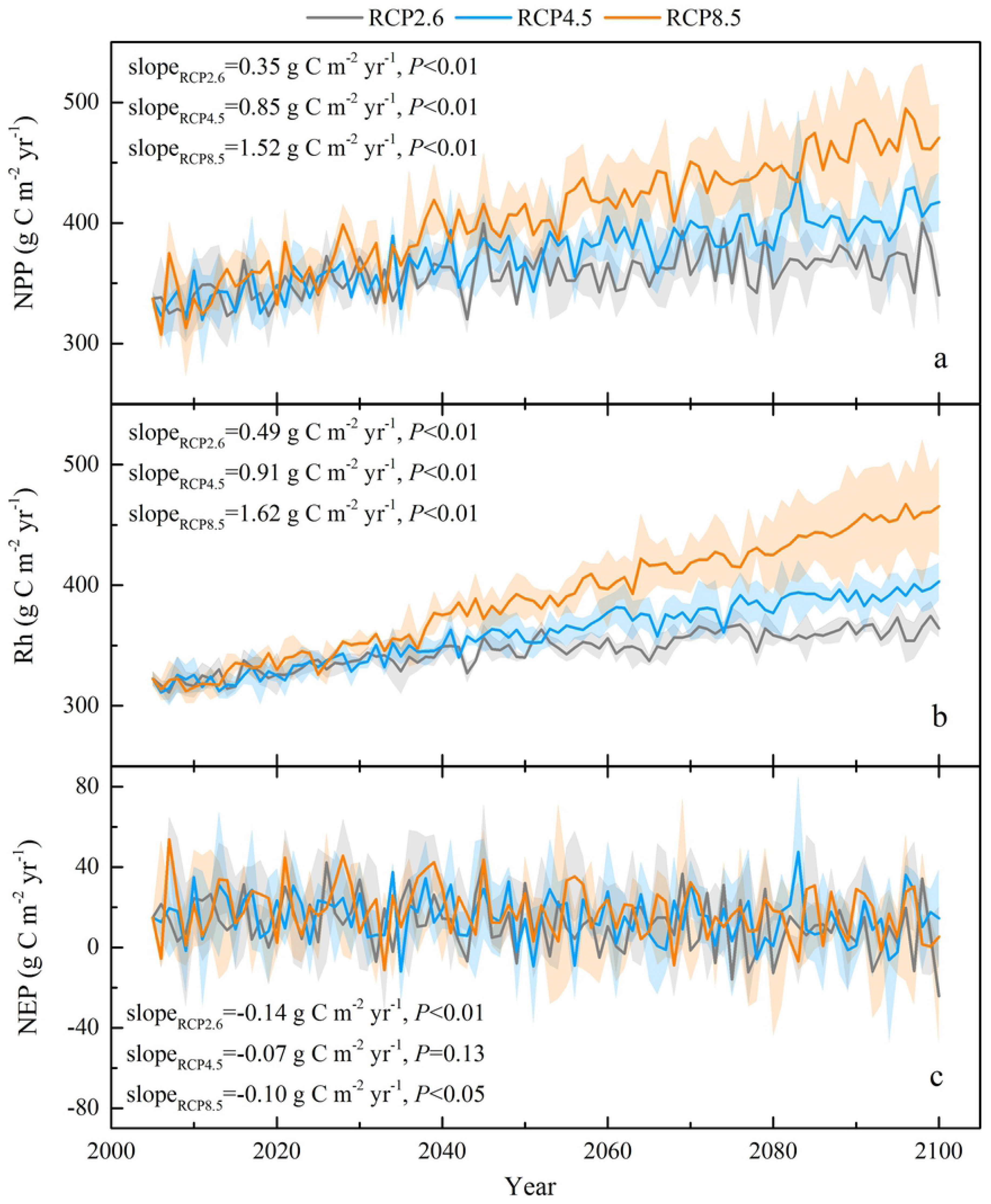
Temporal dynamics of simulated NPP (a), Rh (b), and NEP (c) in the Tibetan Plateau grassland from 2006 to 2100. Solid lines are the three climate models means under different RCP scenarios, and the shading area denotes one standard deviation.

### 3.2 The response of the interannual carbon budget to future climate variability

The simulated carbon fluxes exhibited a positive correlation with precipitation across the three RCP scenarios with the partial correlation coefficients (R) in the range of 0.07 ~ 0.48 (Fig 3). The positive response of the carbon fluxes to annual precipitation variation became stronger with increasing temperature. Higher correlations between carbon fluxes (NPP and Rh) and precipitation were found in RCP8.5 with R values of 0.33 (*P* < 0.05, Fig 3c) and 0.48 (*P* < 0.05, Fig 3f) than those in RCP2.6 (R = 0.25 and 0.34, *P* < 0.05, Fig 3a,d), respectively. Spatially, the carbon fluxes showed a strong positive relationship with precipitation in the midwestern and northeastern part of the Tibetan Plateau grassland (R > 0.3, *P* < 0.05, Fig 3). However, an obvious negative correlation between NEP and precipitation was found in the southern and middle part of the Plateau (Fig 3g-i). This was probably due to the fact that precipitation gradually decreased from the southeast to the northwest [20]. Increased precipitation in the southeastern mountainous area could lead to more cloud cover, and thus have negative effects on photosynthesis. The Rh tended to be most sensitive to the precipitation variation (R:0.34 ~ 0.48, *P* < 0.05, Fig 3d-f), followed by NPP (R:0.25 ~ 0.33, *P* < 0.05, Fig 3a-c) and NEP (R:0.07 ~ 0.08, *P* > 0.05, Fig 3g-i).

**Fig 3.**
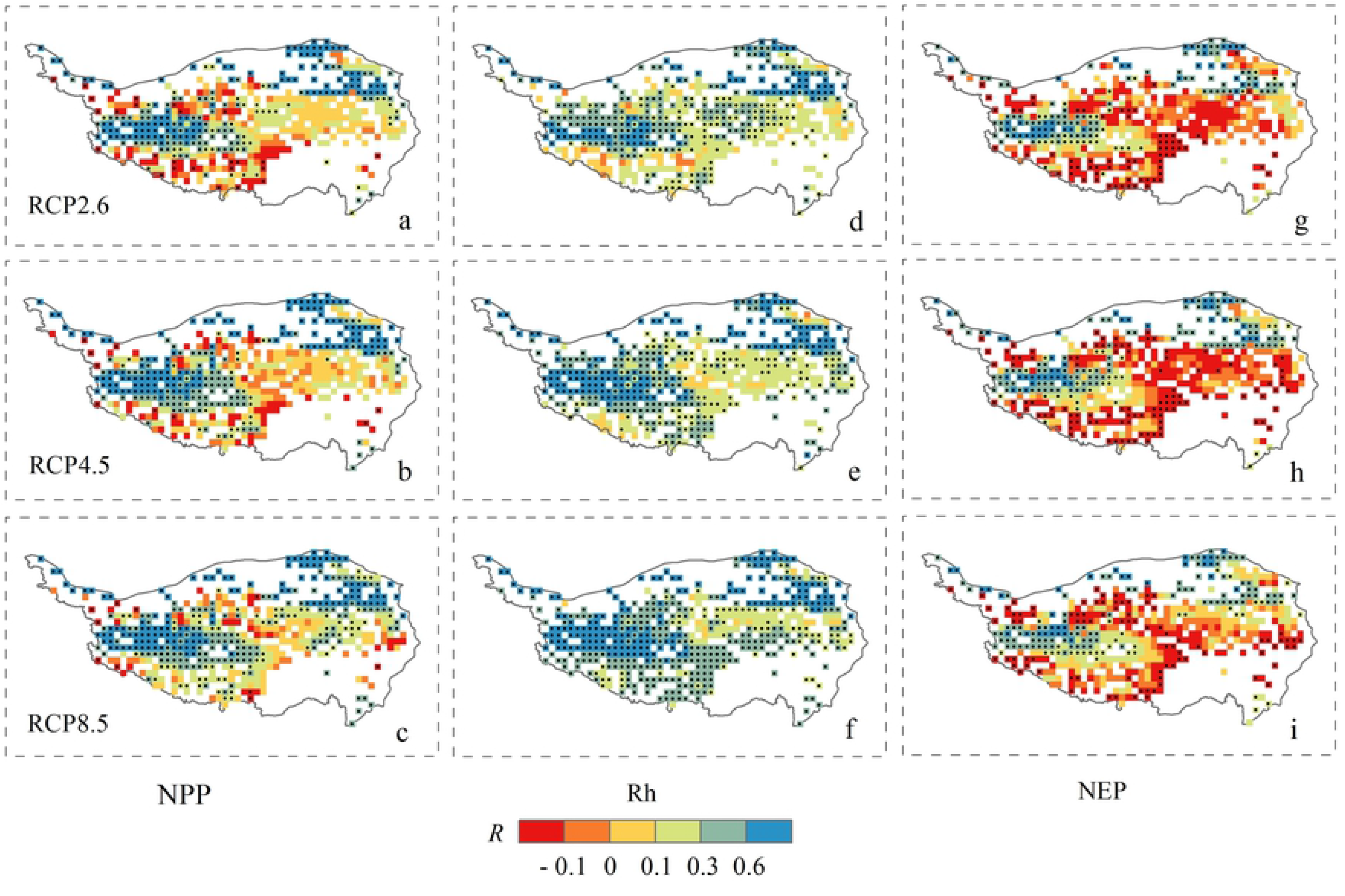
Spatial distributions of the partial correlation coefficients (R) between precipitation and simulated NPP (a, b, c), Rh (d, e, f), and NEP (g, h, i) in the Tibetan Plateau grassland from 2006 to 2100 under the three RCP scenarios. Black point signs showed significance level at P < 0.05.

As illustrated in Fig 4, the simulated carbon fluxes exhibited a positive correlation with temperature (Tmean) (R: 0.03 ~ 0.76), while the R between NEP and Tmean equal to – 0.09. The positive responses of the NPP and Rh to Tmean were stronger in RCP8.5 with R values of 0.54 (*P* < 0.05, Fig 4c) and 0.76 (*P* < 0.05, Fig 4f) than those in RCP2.6 (R = 0.32 and 0.30, *P* < 0.05, Fig 4a,d). Spatially, the strong positive responses of simulated NPP and Rh to Tmean change were found across the majority of the Plateau (R > 0.4, *P* < 0.05, Fig 4a-f), while the NPP and NEP showed an obvious negative responses to increasing temperature in the midwestern and northeastern part of the Plateau (R < −0.2, *P* < 0.05, Fig 4a-c,g-i). This was probably related to the the fact that limited rainfall in these areas could not meet the increase in water demand under warming. The magnitude of Rh increase in response to Tmean (R:0.30 ~ 0.76, *P* < 0.05, Fig 4d-f) was larger than that of NPP (R:0.33 ~ 0.55, *P* < 0.05, Fig 4a-c) and NEP (R:-0.09 ~ 0.14, *P* > 0.05, Fig 4g-i), indicating that warming stimulates respiration more than plant growth and subsequently decreases carbon sink.

**Fig 4.**
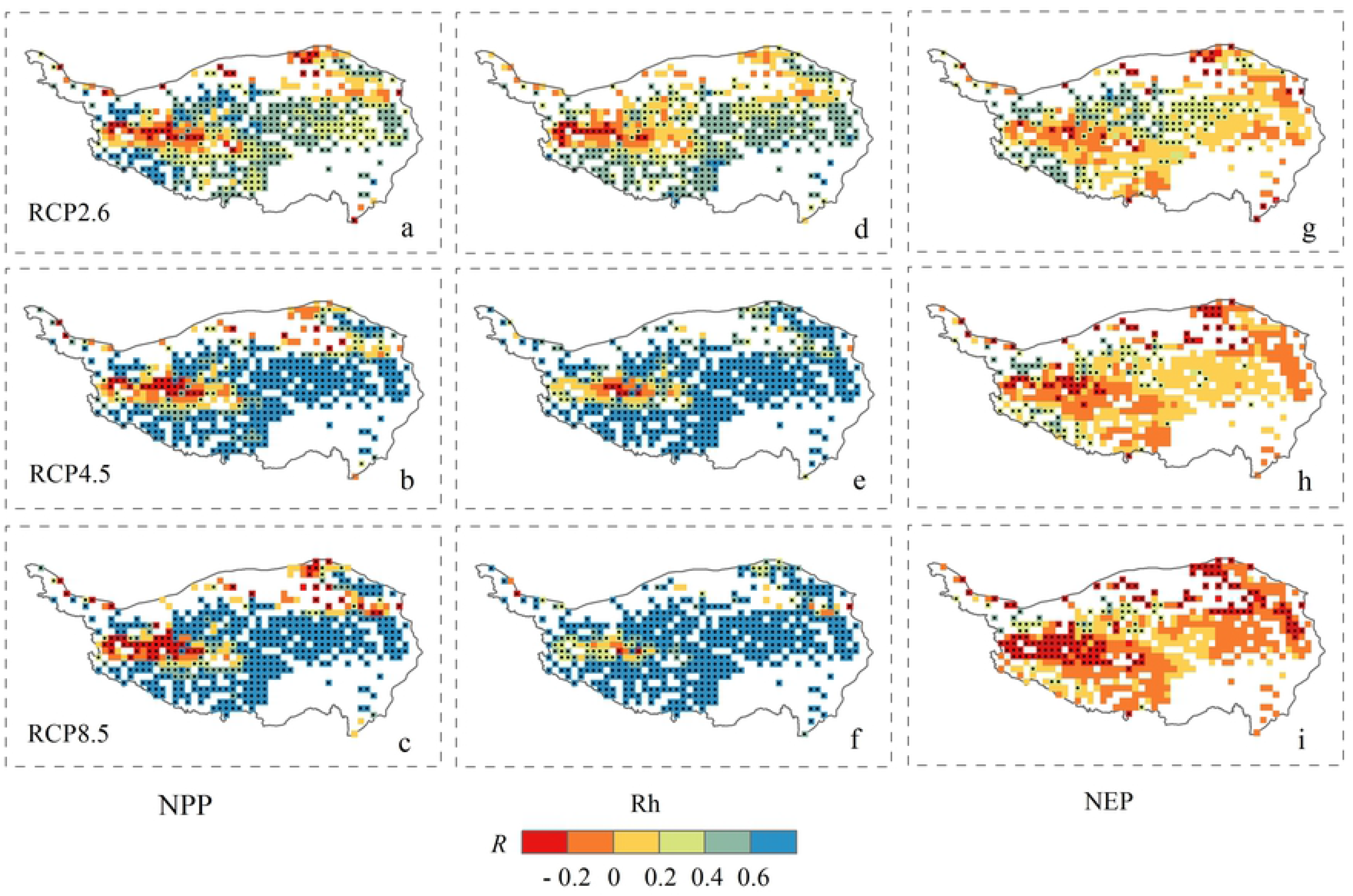
Spatial distributions of the partial correlation coefficients (R) between Tmean and simulated NPP (a, b, c), Rh (d, e, f), and NEP (g, h, i) in the Tibetan Plateau grassland from 2006 to 2100 under the three RCP scenarios. Black point signs showed significance level at P < 0.05.

### 3.3 The response of the decadal carbon budget to future climate change

To quantify the carbon cycle of the Tibetan Plateau grasslands in response to the climate variability, we further evaluate the decadal-scale dynamics of climate and carbon budgets during three time periods (Fig 5). The 2010s ~ 2030s time period was calculated as the difference between the decadal averages of the years 2010 ~ 2019 and 2030 ~ 2039, and the same was done for the 2040s ~ 2060s and 2070s ~ 2090s time periods. The decadal change of carbon fluxes (e.g. ΔNPP and ΔRh) were positively linearly correlated with the ΔTmax (Fig 5d,e) and ΔTmin (Fig 5g,h), with R^2^ in a range of 0.59 ~ 0.67 (*P* < 0.01). It was suggested that the increase in temperature could stimulate both vegetation production and also Rh. Furthermore, the linear slopes indicated that 1% increase in Tmax and Tmin caused a larger increase of Rh by 0.27% (Fig 5c) and 0.24% (Fig 5h) than those of NPP by 0.23% (Fig 5d) and 0.21% (Fig 5g). This suggested that Rh was more sensitive than NPP to increasing temperature. Consequently, the simulated NEP exhibited a decreasing trend under a warming climate. With respect to the response to the decadal change of precipitation, the dynamics of the ΔNPP and ΔRh showed an increasing trend with ΔPrecipitation, while these positive responses tended to be stagnant (Fig 5a,b). As the decadal change of precipitation varied within comparatively small ranges (~ 5%), temperature might play an important role in modulating the carbon fluxes of alpine grassland. Taken together, the changes in ΔNEP suggested that the alpine grassland still behaved as a carbon sink but its capacity declined over the 21st century.

**Fig 5.**
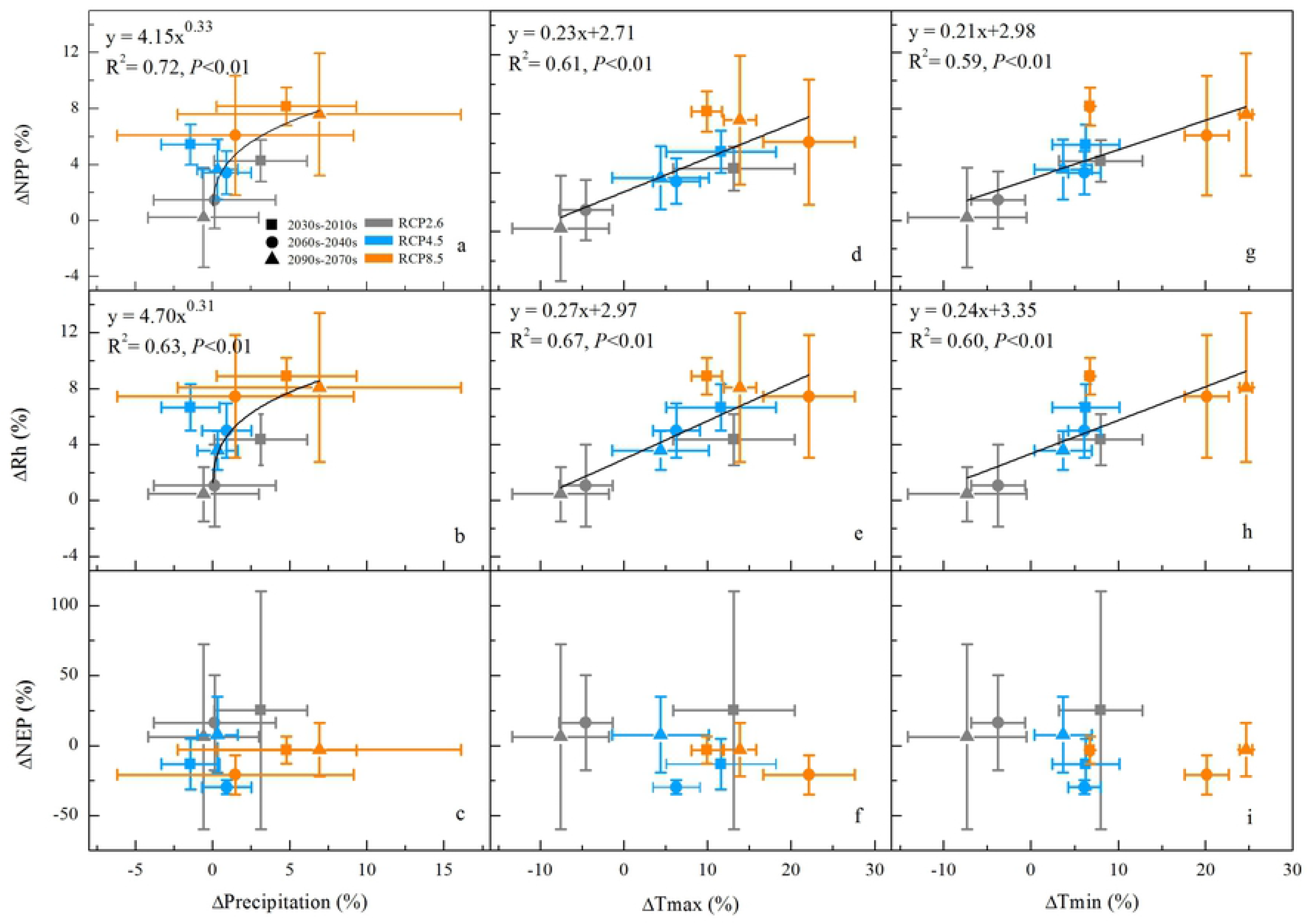
Responses of simulated ΔNPP (a, d, g), ΔRh (b, e, h), and ΔNEP (c, f, i) in the Tibetan Plateau grassland to climate change over the 2010s ~ 2030s, 2040s ~ 2060s and 2070s ~ 2090s time periods under the three RCP scenarios.

Under RCP2.6, the ΔPrecipitation was 3%, 0%, and −1% (Fig 5a-c), with the ΔTmax being 13%, −5%, and −8% (Fig 5d-f), and the ΔTmin 8%, −4%, and −7% (Fig 5g-i) for 2010s ~ 2030s, 2040s ~ 2060s and 2070s ~ 2090s time periods, respectively. This decadal climate change led to a corresponding decrease in ΔNPP from 4% to 1% and 0% for the three time periods (Fig 5a, d, g), and the same decrease in ΔRh (Fig 5b, e and h), and consequently led to a decrease in ΔNEP from 25% to 16% and 6% (Fig 5c, f and i). In response to the continual decrease in temperature and precipitation, the changes in the decadal carbon dynamics were predicted to show a corresponding downward trend in the RCP2.6 scenario. For RCP4.5, the ΔPrecipitation was −1%, 1%, and 0%, with the ΔTmax 12%, 6%, and 4%, and the ΔTmin 6%, 6%, and 4% in the 2010s ~ 2030s, 2040s ~ 2060s and 2070s ~ 2090s time periods, respectively. In this scenario, the reduction in the magnitude of the warming trend was accompanied by stable precipitation, which led to a corresponding decrease in ΔNPP of 5%, 3%, and 4%, and in ΔRh of 7%, 5%, and 4%, respectively. The discrepant magnitudes of the increase in NPP and Rh across different periods, resulting in the variations of the ΔNEP were −13%, −30%, and 8%, respectively. The decrease in ΔNEP in the first two periods was mainly due to a larger increase in Rh than NPP. Under RCP8.5, both the precipitation and temperature were projected to increase when the ΔPrecipitation was 5%, 1%, and 7%, and the ΔTmax was 10%, 22%, and 14%, and the ΔTmin was 7%, 20%, and 25% for 2010s ~ 2030s, 2040s ~ 2060s and 2070s ~ 2090s time periods, respectively. The projected large warming trend over the Tibetan Plateau would favor the growth of alpine grass, with a ΔNPP of 8%, 6%, and 8%, and could also exacerbate carbon decomposition with a ΔRh of 9%, 7%, and 8%, respectively. Consequently, the NEP correspondingly decreased by −3%, −21%, and −3%, respectively. The substantial decrease in the ΔNEP over the 2040s ~ 2060s time period was also partially due to the water stress under strong warming conditions. The dynamics of NEP were largely dependent on the different responses of NPP and Rh to climate change.

### 3.4 The response of the carbon budget to future elevated CO_2_ concentrations

To quantify the effect of increased CO_2_ on the carbon fluxes on alpine grasslands, we conducted two model simulations: with and without CO_2_ fertilization. Then, the difference between the two simulated results was analyzed to evaluate the impacts of CO_2_ on the carbon budget under the three RCP scenarios (Fig 6). Across all scenarios, the increasing CO_2_ concentration accounted for 1.9%, 3.6%, and 7.3% of the increase in NPP (Fig 6a), and 1.7%, 3.0%, and 6.1% of the increase in Rh (Fig 6b) under RCP2.6, RCP4.5, and RCP8.5, respectively. Furthermore, the linear slopes indicated that 1 ppm increase in CO_2_ concentration caused a slightly larger increase of NPP by 0.06 ~ 0.45 g C m^-2^ yr^-1^ (Fig 6a) than those of Rh by 0.06 ~ 0.43 g C m^-2^ yr^-1^ (Fig 6b). Notably, the CO_2_ fertilization effect would be substantially amplified when accompanied by an increase in temperature (slope_Tmax_ = 0.6 °C decade^-1^ and slope_Tmin_ = 0.7 °C decade^-1^, Fig 5d, g) and precipitation (slope = 10.0 mm decade^-1^, Fig 5a) under RCP8.5. By the end of the 21st century (2090s), the ΔNPP and ΔRh both largely increased from RCP2.6 to RCP8.5, ranging from 2.4% to 11.0%. However, the magnitude of the difference in NEP was relatively small, with multi-model mean values of 0.9, 2.3, and 4.9 g C m^-2^ yr^-1^ for RCP2.6, RCP4.5, and RCP8.5, respectively (Fig 6c). The elevated CO_2_ contributed more to plant growth than decomposition, indicating that the Tibetan Plateau grassland was able to sequester carbon from the atmosphere due to CO_2_ fertilization.

**Fig 6.**
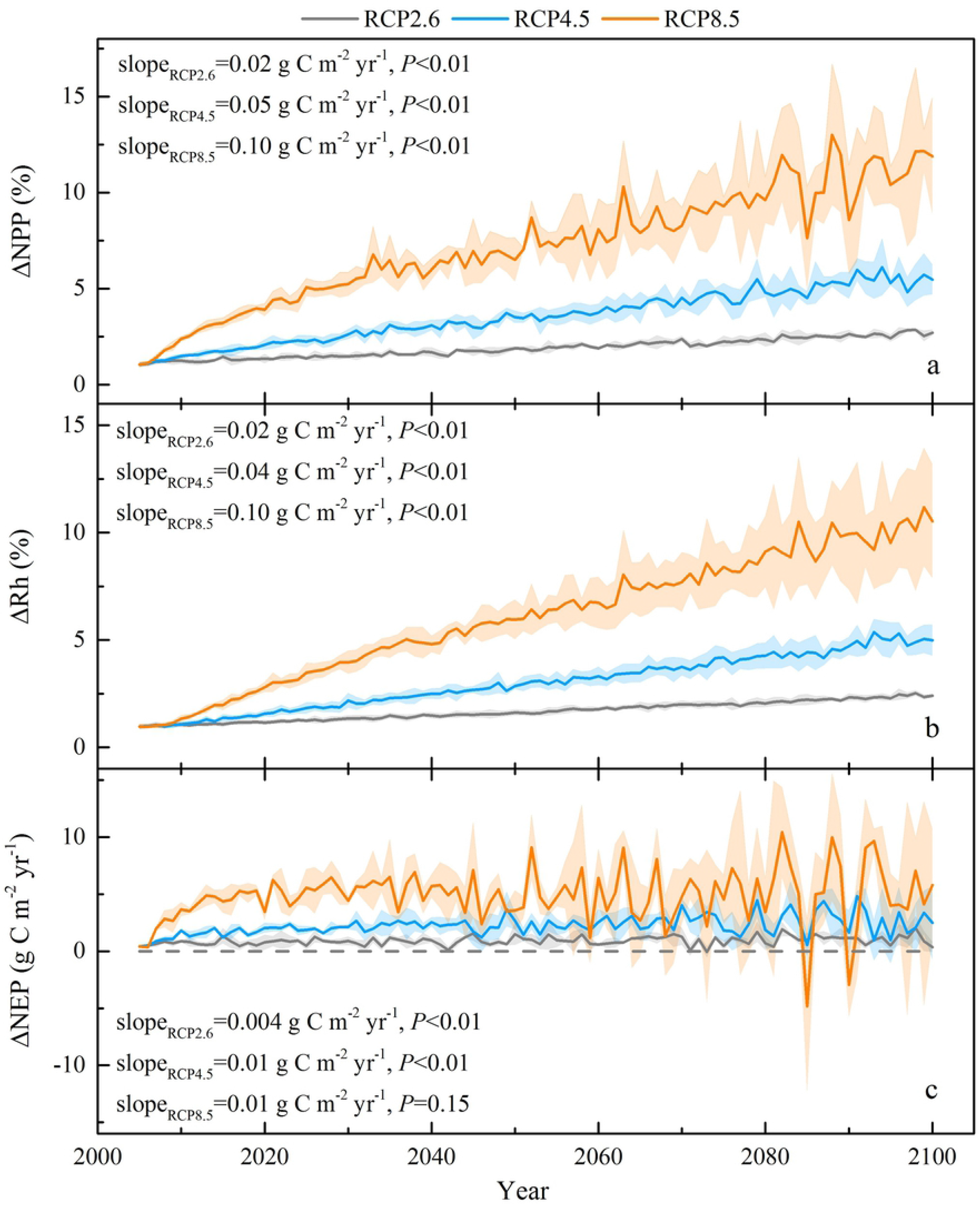
The differences between simulated NPP (a), Rh (b), and NEP (c) with and without CO_2_ effects in the Tibetan Plateau grassland under the three RCP scenarios.

## 4. Discussion

The carbon fluxes of the alpine grassland in the Tibetan Plateau were stimulated to be strongly influenced by the climate change and elevated CO_2_ concentrations [3,5]. Over the 21st century, the Tibetan Plateau grassland NPP and Rh were projected to significantly increase, at rates of 0.35 ~ 1.52 g C m^-2^ yr^-1^ and 0.49 ~ 1.62 g C m^-2^ yr^-1^, respectively, under the three scenarios. These increases in NPP and Rh were primarily due to the effects of warming and elevated CO_2_ concentrations, while the positive impact of precipitation became stable (Fig 5,6). The enhanced vegetation production in future projections was consistent with the model estimations of Gao et al. [7] and Jin et al. [30]. The dynmaics of decadal carbon fluxes (ΔNPP and ΔRh) were predicted to decrease with precipitation and temperature under RCP2.6, and with temperature in a relatively stable precipitation scenario under RCP4.5 (Fig 5). Across all scenarios, the partial correlation (R) between the carbon fluxes (NPP and Rh) and Tmean (Fig 4a-f) were higher than those with precipitation (Fig 3a-f), except for the Rh in RCP4.5. These results suggested that the carbon cycle in the alpine region may be primarily controlled by the temperature increase. However, the partial correlation between carbon sink (NEP) and Tmean was predicted to decrease from 0.14 in RCP2.6 to −0.09 in RCP8.5 under a warmer scenario (Fig 4g,i). The partial correlation of NEP with precipitation were equal to 0.07 and 0.08 in RCP4.5 and RCP8.5 (Fig 3h,i), which were higher than with Tmean (0.04 and −0.09, Fig 4h,i). Furthermore, the decadal carbon fluxes were predicted to decrease with decreasing precipitation, albeit increasing temperature under RCP8.5 during the 2040s ~ 2060s (Fig 5). The role of precipitation was becoming increasingly important in carbon uptake of the Tibetan Plateau grassland. The seasonal dynamics of carbon exchange in alpine grassland under warming were likely to be modulated by water availability [36]. Similar results were found in Inner Mongolia [52], the central and eastern United States [53], and California [54], which indicated that grass production would largely decrease under warmer and drier climate scenarios. The warming accelerated the evapotranspiration and then changed the soil water availability [55,56]. When the soil water availability became limited, the vegetation production might decrease due to suppressed photosynthesis [57] and also reduce root biomass [58]. Shen et al. [6] and Shen et al. [59] further indicated that the precipitation may regulate the response of vegetation phenology and soil respiration to climatic warming on the Tibetan Plateau. The water availability became an increasingly important factor on carbon dynamics on the Tibetan Plateau grasslands, particularly under a warmer climate scenario. These results highlight the significance of accurate projection of precipitation (i.e. the amounts and distributions) under three RCP scenarios, which should be considered in estimates of the carbon fluxes.

The Tibetan Plateau grasslands were projected to be a net carbon sink (16 ~ 25 Tg C yr^-1^) over the 21st century, which was comparable to the results of Jin et al. [30] with NEP ranging from 18 to 46 Tg C yr^-1^. However, the temporal dynamics of NEP all exhibited a slightly decreasing trend, with slopes of −0.07 ~ −0.14 g C m^-2^ yr^-1^ across the three RCP scenarios. This was probably due to the fact that warming induced more carbon inputs into the soil and accelerated respiration rates (Rh), leading to reduced carbon uptake [60,61]. On the other side, Cao and Woodward [62] found that the increasing trend in carbon uptake would be diminished by the acclimated response to climate variability and CO_2_ fertilization effect. Consequently, the temporal dynamics of simulated SOC increased 10 ~ 15 Mg C ha ^-1^ (i.e., 1.4 ~ 2.1 Pg C for 1.4 × 10^6^ km^2^) over the 21st century, while the changing rate showed a decreasing trend of −0.12 ~ −0.06 g C m^-2^ yr^-1^ (*P*<0.01) under the three RCP scenarios. These results were in line with the projections from the CMIP5 [63], but inconsistent with the findings of Crowther et al. [64], due largely to the high initial SOC density in high-latitude areas. Our findings suggested that the carbon sequestration in the vegetation and soil was projected to increase under three RCP scenarios, but the carbon sequestration rate was predicted to decrease gradually on the Tibetan Plateau grasslands.

The carbon dynamics exhibited a large discrepancy in response to climate change with and without the effect of elevated CO_2_ concentrations. The carbon uptake by vegetation could be strongly stimulated by CO_2_ fertilization, especially in July [65]. The increasing CO_2_ concentration accounted for 1.9% ~ 7.3% of the increase in NPP (Fig 6a), which was less than the range of 9% ~ 11% in the perennial grassland experiment at the Cedar Creek Ecosystem Science Reserve in central Minnesota during 1998 ~ 2010 [66]. Furthermore, the slope of NPP was predicted to increase by 18% ~ 29% under CO_2_ fertilization among all scenarios, which was slightly less than the estimates of Piao et al. [3] (~ 39%) on the Tibetan Plateau grasslands during the past five decades. This was probably due to the effect of CO_2_ fertilization, which tended to be saturated over time, and the discrepancy that existed in different models. Furthermore, the response of Rh to CO_2_ was of the same magnitude (1.7% ~ 6.1%) as that in NPP during the 21st century. The elevated CO_2_ concentrations generally had a positive effect on soil respiration by regulating the carbon uptake and allocation, and on the availability of substrate and soil water [67]. Consequently, the almost consistent increase in NPP and Rh caused by the elevated CO_2_ on the alpine grassland resulted in a relatively small difference in NEP (0.9 ~ 4.9 g C m^-2^ yr^-1^, Fig 6c). However, the grassland carbon cycle was projected to be amplified by around two to three times due to higher CO_2_ concentrations under RCP8.5 compared to that in RCP2.6 and RCP4.5. The different potential responses to elevated CO_2_ levels were partly due to the different climate conditions among the three RCP scenarios. For RCP2.6, the increasing CO_2_ associated with a decreasing trend in temperature and precipitation had a minor effect on the carbon cycle (Fig 6a). Furthermore, the elevated CO_2_ favored plant growth limitedly in RCP4.5, as the continually warming and stable precipitation may induce water stress (Fig 6b). However, increasing temperature and precipitation, together with increasing CO_2_, greatly enhanced the carbon uptake in the alpine ecosystem under RCP8.5 (Fig 6c).

Furthermore, the dynamically downscaled climate data using regional climate models (RCMs) indicated an improved performance compared to the coarse output from global climate models (GCMs) [51,68]. Jin et al. [30] indicated that the increased resolution of RCMs was preferable for capturing the regional climate patterns, especially under the complex surface characteristics of the Tibetan Plateau. There are also substantial uncertainties in projected climate change for RCMs, coming from the driving GCMs and human action [69,70]. Moreover, without taking the effect of nitrogen (N) into account, the quantitative attribution of future climate change and elevated CO_2_ concentrations on the carbon dynamics of the Tibetan Plateau grasslands should be considered merely suggestive. The N is regarded as a limiting factor, controlling the carbon uptake and further influencing the climate – carbon cycle feedback [71,72]. However, most of the current carbon cycle models lack N dynamics [71]. Further efforts should be made to incorporate N feedback to constrain the uncertainty of the carbon cycle in response to a changing environment [73,74].

## 5. Conclusions

By applying a process-based biogeochemistry model, CENTURY, we quantified the carbon dynamics on the Tibetan Plateau grasslands in response to climate change and elevated CO_2_ under the RCP scenarios. The Tibetan Plateau grasslands behaved as a net carbon sink with the NEP of 16 ~ 25 Tg C yr^-1^ over the 21st century. However, the potential capacity for carbon sequestration on the Tibetan Plateau grasslands was predicted to decrease gradually with the slopes ranging from −0.14 to −0.07. The partial correlation between carbon fluxes and climate indictated that warming stimulated respiration more than plant growth. However, the elevated CO_2_ contributed more to plant growth than decomposition, which could offset the warming-induced carbon loss. The interannual and decadal-scale dynamics of the carbon fluxes were primarily controlled by temperature, while the role of precipitation became increasingly important in modulating carbon budgets in the alpine grassland. The stable precipitation accompanied by noticeable warming in RCP4.5 (2010s ~ 2030s and 2040s ~ 2060s) and RCP8.5 (2040s ~ 2060s) led to a reduction of carbon sink (ΔNEP) by −13%, −30% and −21%, respectively. These results highlighted the importance of precipitation in regulating the contribution of CO_2_ fertilization and warming effect on carbon dynamics in a warmer climate scenario.

## Acknowledgments

This work was supported by National Key R&D Program of China (No. 2017YFB0504000) and the Postdoctoral Science Foundation of LASG Dean (No. 7-091162) to P.F.H‥ We are grateful to the Chinese Ecosystem Research Network (CERN) and the Botanical Division of Chinese Biodiversity Information Center, Institute of Botany for kindly providing the Haibei observation datasets.

## References

1. Chen L, Liang J, Qin S, Liu L, Fang K, et al. (2016) Determinants of carbon release from the active layer and permafrost deposits on the Tibetan Plateau. Nature Communications 7.

2. Yang K, Wu H, Qin J, Lin C, Tang W, et al. (2014) Recent climate changes over the Tibetan Plateau and their impacts on energy and water cycle: A review. Global and Planetary Change 112: 79–91.

3. Piao S, Tan K, Nan H, Ciais P, Fang J, et al. (2012) Impacts of climate and CO_2_ changes on the vegetation growth and carbon balance of Qinghai-Tibetan grasslands over the past five decades. Global and Planetary Changes 98–99: 73–80.

4. Ding J, Chen L, Ji C, Hugelius G, Li Y, et al. (2017) Decadal soil carbon accumulation across Tibetan permafrost regions. Nature Geoscience 10: 420–424.

5. Zhuang Q, He J, Lu Y, Ji L, Xiao J, et al. (2010) Carbon dynamics of terrestrial ecosystems on the Tibetan Plateau during the 20th century: An analysis with a process-based biogeochemical model. Global Ecology and Biogeography 19: 649–662.

6. Shen M, Zhang G, Cong N, Wang S, Kong W, et al. (2014) Increasing altitudinal gradient of spring vegetation phenology during the last decade on the Qinghai-Tibetan Plateau. Agricultural and Forest Meteorology s 189–190: 71–80.

7. Gao Q, Guo Y, Xu H, Ganjurjav H, Li Y, et al. (2016) Climate change and its impacts on vegetation distribution and net primary productivity of the alpine ecosystem in the Qinghai-Tibetan Plateau. Science of the Total Environment 554–555: 34–41.

8. Friend AD, Lucht W, Rademacher TT, Keribin R, Betts R, et al. (2014) Carbon residence time dominates uncertainty in terrestrial vegetation responses to future climate and atmospheric CO_2_. Proceedings of the National Academy of Sciences of the United States of America 111: 3280–3285.

9. Kicklighter DW, Cai Y, Zhuang Q, Parfenova EI, Paltsev S, et al. (2014) Potential influence of climate-induced vegetation shifts on future land use and associated land carbon fluxes in Northern Eurasia. Environmental Research Letters 9: 156–166.

10. Dieleman CM, Branfireun BA, Lindo Z (2017) Northern peatland carbon dynamics driven by plant growth form—the role of graminoids. Plant and Soil 415: 25–35.

11. Morales P, Hickler T, Rowell DP, Smith B, Sykes MT (2007) Changes in European ecosystem productivity and carbon balance driven by regional climate model output. Global Change Biology 13: 108–122.

12. Koven CD, Ringeval B, Friedlingstein P, Ciais P, Cadule P, et al. (2011) Permafrost carbon-climate feedbacks accelerate global warming. Proceedings of the National Academy of Sciences of the United States of America 108: 14769–14774.

13. Schaefer K, Lantuit H, Romanovsky VE, Schuur EAG, Witt R (2014) The impact of the permafrost carbon feedback on global climate. Environmental Research Letters 9.

14. Schädel C, Bader MKF, Schuur EAG, Biasi C, Bracho R, et al. (2016) Potential carbon emissions dominated by carbon dioxide from thawed permafrost soils. Nature Climate Change 6.

15. Schaphoff S, Heyder U, Ostberg S, Gerten D, Heinke J, et al. (2013) Contribution of permafrost soils to the global carbon budget. Environmental Research Letters 8: 3865–3879.

16. Morgan JA, Lecain DR, Mosier AR, Milchunas DG (2001) Elevated CO_2_ enhances water relations and productivity and affects gas exchange in C3 and C4 grasses of the Colorado shortgrass steppe. Global Change Biology 7: 451–466.

17. Donohue RJ, Roderick ML, Mcvicar TR, Farquhar GD (2013) Impact of CO_2_ fertilization on maximum foliage cover across the globe’s warm, arid environments. Geophysical Research Letters 40: 3031–3035.

18. Chen H, Zhu Q, Peng C, Wu N, Wang Y, et al. (2013) The impacts of climate change and human activities on biogeochemical cycles on the Qinghai-Tibetan Plateau. Global change biology 19: 2940–2955.

19. Liu XD, Chen BD (2000) Climatic warming in the Tibetan plateau during recent decades. International Journal of Climatology 20: 1729–1742.

20. Kang SC, Xu YW, You QL, Flügel WA, Pepin N, et al. (2010) Review of climate and cryospheric change in the Tibetan Plateau. Environmental Research Letters 5: 75–82.

21. Su F, Duan X, Chen D, Hao Z, Cuo L (2013) Evaluation of the Global Climate Models in the CMIP5 over the Tibetan Plateau. Journal of Climate 26: 3187–3208.

22. Tan K, Ciais P, Piao S, Wu X, Tang Y, et al. (2010) Application of the ORCHIDEE global vegetation model to evaluate biomass and soil carbon stocks of Qinghai-Tibetan grasslands. Global Biogeochemical Cycles 24.

23. Lin X, Han P, Zhang W, Wang G (2017) Sensitivity of alpine grassland carbon balance to interannual variability in climate and atmospheric CO_2_ on the Tibetan Plateau during the last century. Global and Planetary Change 154: 23–32.

24. Saito M, Kato T, Tang Y (2009) Temperature controls ecosystem CO_2_ exchange of an alpine meadow on the northeastern Tibetan Plateau. Global Change Biology 15: 221–228.

25. Kato T, Tang Y, Gu S, Hirota M, Du M, et al. (2006) Temperature and biomass influences on interannual changes in CO_2_ exchange in an alpine meadow on the Qinghai-Tibetan Plateau. Global Change Biology 12: 1285–1298.

26. Piao S, Cui M, Chen A, Wang X, Ciais P, et al. (2011) Altitude and temperature dependence of change in the spring vegetation green-up date from 1982 to 2006 in the Qinghai-Xizang Plateau. Agricultural and Forest Meteorology 151: 1599–1608.

27. Yu H, Luedeling E, Xu J (2010) Winter and spring warming result in delayed spring phenology on the Tibetan Plateau. Proceedings of the National Academy of Sciences of the United States of America 107: 22151–22156.

28. Zhang G, Zhang Y, Dong J, Xiao X (2013) Green-up dates in the Tibetan Plateau have continuously advanced from 1982 to 2011. Proceedings of the National Academy of Sciences of the United States of America 110: 4309–4314.

29. Gao Y, Zhou X, Wang Q, Wang C, Zhan Z, et al. (2013) Vegetation net primary productivity and its response to climate change during 2001-2008 in the Tibetan Plateau. Science of the Total Environment 444: 356–362.

30. Jin Z, Zhuang Q, He JS, Zhu X, Song W (2015) Net exchanges of methane and carbon dioxide on the Qinghai-Tibetan Plateau from 1979 to 2100. Environmental Research Letters 10.

31. Klein JA, Harte J, Zhao X-Q (2007) Experimental warming, not grazing, decreases rangeland quality on the Tibetan Plateau. Ecological Applications 17: 541–557.

32. Li N, Genxu W, Yan Y, Yongheng G, Guangsheng L (2011) Plant production, and carbon and nitrogen source pools, are strongly intensified by experimental warming in alpine ecosystems in the Qinghai-Tibet Plateau. Soil Biology and Biochemistry 43: 942–953.

33. Fu G, Zhang X, Zhang Y, Shi P, Li Y, et al. (2013) Experimental warming does not enhance gross primary production and above-ground biomass in the alpine meadow of Tibet. Journal of Applied Remote Sensing 7.

34. Ganjurjav H, Gao Q, Gornish ES, Schwartz MW, Liang Y, et al. (2016) Differential response of alpine steppe and alpine meadow to climate warming in the central Qinghai-Tibetan Plateau. Agricultural and Forest Meteorology 223: 233–240.

35. Piao S, Fang J, Ciais P, Peylin P, Huang Y, et al. (2009) The carbon balance of terrestrial ecosystems in China. Nature 458: 1009–1013.

36. Zhu J, Zhang Y, Jiang L (2017) Experimental warming drives a seasonal shift of ecosystem carbon exchange in Tibetan alpine meadow. Agricultural and Forest Meteorology: 242–249.

37. Taylor KE, Stouffer RJ, Meehl GA (2012) An overview of CMIP5 and the experiment design. Bulletin of the American Meteorological Society 93: 485–498.

38. Moss RH, Edmonds JA, Hibbard KA, Manning MR, Rose SK, et al. (2010) The next generation of scenarios for climate change research and assessment. Nature 463: 747–756.

39. Parton WJ, Schimel DS, Cole C, Ojima D (1987) Analysis of factors controlling soil organic matter levels in Great Plains grasslands. Soil Science Society of America Journal 51: 1173–1179.

40. Hall D, Ojima D, Parton W, Scurlock J (1995) Response of temperate and tropical grasslands to CO_2_ and climate change. Journal of Biogeography; 537–547.

41. Parton W, Scurlock J, Ojima D, Gilmanov T, Scholes R, et al. (1993) Observations and modeling of biomass and soil organic matter dynamics for the grassland biome worldwide. Global biogeochemical cycles 7: 785–809.

42. Parton W, Scurlock J, Ojima D, Schimel D, Hall D (1995) Impact of climate change on grassland production and soil carbon worldwide. Global Change Biology 1: 13–22.

43. Chimner RA, Cooper DJ, Parton WJ (2002) Modeling carbon accumulation in Rocky Mountain fens. Wetlands 22: 100–110.

44. McGuire A, Melillo J, Randerson J, Parton W, Heimann M, et al. (2000) Modeling the effects of snowpack on heterotrophic respiration across northern temperate and high latitude regions: Comparison with measurements of atmospheric carbon dioxide in high latitudes. Biogeochemistry 48: 91–114.

45. Zhu X, Yu G, Wang Q, Gao Y, Zhao X, et al. (2013) The interaction between components of ecosystem respiration in typical forest and grassland ecosystems. Acta Ecologica Sinica 33: 6925–6934.

46. Zhang Y, Wei Q, Zhou C, Ding M, Liu L, et al. (2014) Spatial and temporal variability in the net primary production of alpine grassland on the Tibetan Plateau since 1982. Journal of Geographical Sciences 24: 269–287.

47. Pan S, Dangal SR, Tao B, Yang J, Tian H (2015) Recent patterns of terrestrial net primary production in Africa influenced by multiple environmental changes. Ecosystem Health and Sustainability 1: 1–15.

48. Tian H, Melillo JM, Kicklighter DW, McGuire AD, Helfrich JV, et al. (1998) Effect of interannual climate variability on carbon storage in Amazonian ecosystems. Nature 396: 664–667.

49. Mitchell TD, Jones PD (2005) An improved method of constructing a database of monthly climate observations and associated high-resolution grids. International journal of climatology 25: 693–712.

50. Harris I, Jones P, Osborn T, Lister D (2014) Updated high-resolution grids of monthly climatic observations-the CRU TS3. 10 Dataset. International Journal of Climatology 34: 623–642.

51. Jones C, Giorgi F, Asrar G (2011) The Coordinated Regional Downscaling Experiment: CORDEX, an international downscaling link to CMIP5. CLIVAR exchanges 56: 34–40.

52. Kang X, Hao Y, Li C, Cui X, Wang J, et al. (2011) Modeling impacts of climate change on carbon dynamics in a steppe ecosystem in Inner Mongolia, China. Journal of Soils and Sediments 11: 562–576.

53. Behrman KD, Kiniry JR, Winchell M, Juenger TE, Keitt TH (2013) Spatial forecasting of switchgrass productivity under current and future climate change scenarios. Ecological Applications 23: 73–85.

54. Lenihan JM, Bachelet D, Neilson RP, Drapek R (2008) Response of vegetation distribution, ecosystem productivity, and fire to climate change scenarios for California. Climatic Change 87: 215–230.

55. Cook BI, Smerdon JE, Seager R, Coats S (2014) Global warming and 21 st century drying. Climate Dynamics 43: 2607–2627.

56. Trenberth KE, Dai A, Schrier GVD, Jones PD, Barichivich J, et al. (2013) Global warming and changes in drought. Nature Climate Change 4: 17–22.

57. Liu Y, Wang T, Huang M, Yao Y, Ciais P, et al. (2016) Changes in interannual climate sensitivities of terrestrial carbon fluxes during the 21st century predicted by CMIP5 Earth system models. Journal of Geophysical Research Biogeosciences 121.

58. de Vries FT, Brown C, Stevens CJ (2016) Grassland species root response to drought: consequences for soil carbon and nitrogen availability. Plant and Soil 409: 297–312.

59. Shen ZX, Li YL, Fu G (2015) Response of soil respiration to short-term experimental warming and precipitation pulses over the growing season in an alpine meadow on the Northern Tibet. Applied Soil Ecology 90: 35–40.

60. Dorrepaal E, Toet S, Logtestijn RSPV, Swart E, Weg MJVD, et al. (2009) Carbon respiration from subsurface peat accelerated by climate warming in the subarctic. Nature 460: 616–619.

61. Curiel YJ, Baldocchi DD, Gershenson A, Goldstein A, Misson L, et al. (2007) Microbial soil respiration and its dependency on carbon inputs, soil temperature and moisture. Global Change Biology 13: 2018–2035.

62. Cao M, Woodward FI (1998) Dynamic responses of terrestrial ecosystem carbon cycling to global climate change. Nature 393: 249–252.

63. Stocker TF, Qin D, Plattner G, Tignor M, Allen S, et al. (2013) Climate change 2013: the physical science basis. Intergovernmental panel on climate change, working group I Contribution to the IPCC fifth assessment report (AR5). New York.

64. Crowther T, Todd-Brown K, Rowe C, Wieder W, Carey J, et al. (2016) Quantifying global soil carbon losses in response to warming. Nature 540: 104–108.

65. Zeng N, Zhao F, Collatz GJ, Kalnay E, Salawitch RJ, et al. (2014) Agricultural Green Revolution as a driver of increasing atmospheric CO_2_ seasonal amplitude. Nature 515: 394–397.

66. Reich PB, Hobbie SE (2013) Decade-long soil nitrogen constraint on the CO_2_ fertilization of plant biomass. Nature Climate Change 3: 278–282.

67. Wan S, Norby RJ, Ledford J, Weltzin JF (2007) Responses of soil respiration to elevated CO_2_, air warming, and changing soil water availability in a model old-field grassland. Global Change Biology 13: 2411–2424.

68. Giorgi F, Jones C, Asrar GR (2009) Addressing climate information needs at the regional level: the CORDEX framework. Bulletin-World Meteorological Organization: 175–183.

69. Foley AM (2010) Uncertainty in regional climate modelling: A review. Progress in Physical Geography 34: 647–670.

70. Giorgi F, Gutowski WJ (2015) Regional Dynamical Downscaling and the CORDEX Initiative. Annual Review of Environment and Resources 40: 1–24.

71. Wårlind D, Smith B, Hickler T, Arneth A (2014) Nitrogen feedbacks increase future terrestrial ecosystem carbon uptake in an individual-based dynamic vegetation model. Biogeosciences Discussions 11: 6131–6146.

72. Sokolov AP, Kicklighter DW, Melillo JM, Felzer BS, Schlosser CA, et al. (2008) Consequences of Considering Carbon-Nitrogen Interactions on the Feedbacks between Climate and the Terrestrial Carbon Cycle. Journal of Climate 21: 3776–3796.

73. Wieder WR, Cleveland CC, Smith WK, Todd-Brown K (2015) Future productivity and carbon storage limited by terrestrial nutrient availability. Nature Geoscience 8: 441–444.

74. Piao S, Sitch S, Ciais P, Friedlingstein P, Peylin P, et al. (2013) Evaluation of terrestrial carbon cycle models for their response to climate variability and to CO_2_ trends. Global Change Biology 19: 2117–2132.

